# The knockdown of the snoRNA-jouvence blocks the cell proliferation and leads to cell death of human primary cancerous glioblastoma cells

**DOI:** 10.1101/2024.08.23.609357

**Authors:** Lola Jaque-Cabrera, Julia Buggiani, Jérôme Bignon, Patricia Daira, Nathalie Bernoud-Hubac, Jean-René Martin

**Author notes:** corresponding author: Jean-René Martin. Institut de Biologie Intégrative de la Cellule (I2BC) CNRS UMR-9198, 1 avenue de la terrasse, Bâtiment 21 F91198 Gif-sur-Yvette cedex, France.

## Abstract

Cancer research aims to identify new therapeutic targets and provide patients with more effective therapies that generate fewer deleterious side effects. Previous studies have demonstrated the role of small nucleolar RNAs (snoRNAs) in many physiological and pathological cellular processes, including cancers. SnoRNAs are a group of non-coding RNAs involved in different post-transcriptional modifications of ribosomal RNAs. Recently, we have identified a new snoRNA, named *jouvence* (snoRNA-jou) in humans. Here, we characterize the effect of the knockdown of the snoRNA-jouvence by sh-lentivirus on human primary glioblastoma cells. The sh-lentivirus induces a significant decrease of cell proliferation, proportional to the amount of lentivirus used. For example, with a MOI-20, a decrease of more than 50% is observed after only 72 hours, and even up to 80% after 10 days. Moreover, a EdU proliferative test confirms this decrease in cell proliferation, while a TUNEL test reveals the presence of apoptotic cells. A RNA-Seq analysis, validated by RT-qPCR, reveals that the snoRNA-jou knockdown induces a significant deregulation of several key genes involved in the cell cycle regulation, such as cyclins B1, A2 and p21, indicating a cell cycle blockage. Altogether, these results allow hypothesising that the knockdown of the snoRNA-jouvence could potentially be use as a new anti-cancer treatment (sno-Therapy).

## BACKGROUND

Cancer research aims to understand the cellular and molecular mechanisms involved, in order to identify new therapeutic targets and provide patients with more effective therapies that generate fewer side undesirable and toxic effects. Glioblastoma is the most common primary malignant brain tumour, with a worldwide incidence of 10 per 100,000 population [1]. Despite advances in research in surgery, radiotherapy and chemotherapy improving the survival and quality of life of patients, this disease remains for the most cases incurable, with a survival rate of only 14 months after diagnosis [1]. Patients have a high rate of recurrence and resistance to treatment. These figures therefore make it a crucial public health issue.

Recently, studies have demonstrated the role of small nucleolar RNAs (snoRNAs) in many physiological and pathological cellular processes, including cancers. Indeed, some snoRNAs show aberrant expression in several types of cancers and can also be used, for some of them, as a marker of disease progression [2–5]. They are also deregulated between different stages and different types of cancers [6,7]. SnoRNAs are a group of non-coding RNAs contained in the nucleolus, the largest structure in the nucleus of eukaryotic cells involved in the synthesis of ribosomal RNAs (rRNAs) as well as the assembly and maturation of ribosomes [8]. SnoRNAs associate with proteins to form a complex, called small nucleolar ribonucleoprotein, carrying out different post-transcriptional modifications of rRNAs. The snoRNAs are divided into two major classes: C/D box snoRNAs are responsible for 2-O-methylation, while H/ACA box snoRNAs are responsible for the pseudouridylation of target nucleotides. Pseudouridylation, corresponding to the conversion of a uridine into pseudo-uridine, takes place at the level of the internal loops of snoRNAs, called pseudouridylation pockets [8]. Pseudouridine is one of the most abundant post-transcriptional modifications of RNA nucleotides [9]. As well known, the role of ribosomes in protein translation is essential for cell growth and proliferation [10–12]. However, the excessive and uncontrolled proliferation of cancer cells leads to the formation of malignant tumor. It is therefore accepted that the involvement of snoRNAs in the modification of rRNAs could contribute to the progression of cancers [7].

Our recent research has demonstrated the involvement of a new snoRNA identified in Drosophila, called snoRNA-*jouvence* (snoRNA-jou), in process of longevity and protection against neurodegeneration [13,14]. Since snoRNAs are highly conserved during evolution [8], its counterpart in mice and human have been identified, since they were not yet annotated in the genome [13,15]. The human genome has a single copy of snoRNA-jou of 159 pb, located on chromosome 11, in an intron of the TEAD1 gene [13,15]. The TEAD family of transcription factors plays different roles in cell proliferation, as well as tissue regeneration and homeostasis [16]. Structurally, human snoRNA-jou belongs to the class of H/ACA box snoRNAs [15]. However, it has a non-canonical secondary structure, i.e. it does not exhibit a classical “double hairpin” structure like most H/ACA box snoRNAs. However, its structure does not seem to impact its function, since a pseudouridylation pocket has been identified that can likely pseudouridylate 18S rRNA at position 1397 [15]. In order to characterize the role of snoRNA-jou in humans, first, its expression was studied on nine human cell lines, including immortalized cell lines. Three normal (non-cancerous) cell lines were studied; HEK (embryonic kidney), MRC5 (embryonic lung), RPE1 (retinal pigment epithelium), as well as six cancer cell lines : HCT116 and Caco2 (colon cancer), HeLa (cervical cancer), PC3 (prostate cancer), U87 (glioblastoma), A549 (lung cancer). In all these cell types, the snoRNA-jou is expressed at a low level, demonstrating that the snoRNA-jou is expressed in many cell types, either in a pathological context or not [15]. Studies were then carried out to study the overexpression and inactivation (knockdown) of snoRNA-jou in these lines. Overexpression of snoRNA-jou induces an increase in proliferation in all these cell types; while knockdown induces a decrease in cell proliferation [15]. In addition, a transcriptomic analysis (RNA-Seq) performed on the HCT116 cells overexpressing snoRNA-jou indicates the deregulation of several genes, while KEGG and GO analysis have suggested a genomic signature of a de-differentiation process, compatible with a rejuvenation. Inversely, again on the HCT116 cells, a RNA-Seq performed on the knockdown of snoRNA-jou reveals a drastic decrease of the majority of genes encoding for ribosomal proteins, indicating a collapse of the ribogenesis, as well as several genes involved in the splicesome [15]. These two last effects of the knockdown correlates perfectly with the important decrease in cell proliferation phenotype.

Up to now, the snoRNA-jou knockdown experiments were carried out using small interfering RNAs (siRNAs) on immortalized cell lines [15]. However, the transfection of cells by vectors containing siRNAs is transient. This means that the DNA is not incorporated into the genome of the cells and therefore the expression of siRNAs is no longer present in the daughter cells after cell division. Here, we decide to induce snoRNA-jou knockdown using lentiviruses containing short hairpin RNAs (shRNAs) directed against snoRNA-jou. The transduction of shRNAs by a viral vector makes it possible to more easily intervene on primary cells, which are usually more difficult to transfect by other techniques. Lentivirus are RNA viruses that can integrate their vector into the genome of the host cell and therefore allow stable expression of the transgene, here the shRNA. We decide to begin experiments first on the HCT116 whose have been well characterized in our former study [15], in the aim to validate the sh-lentivirus approach, and second on primary cells, the GBM14-CHA cells (xenograft), derived from a patient with glioblastoma. These positive results suggest that the knockdown of snoRNA-jou could make it possible to block the proliferation of cancer cells and therefore could potentially represent a new therapeutic tool against certain cancers.

## RESULTS

### Knockdown of the snoRNA-jou using sh-lentivirus decreases cancerous cell proliferation

To study the snoRNA-jou knockdown effects, human cancerous cells were transduced with sh-lentivirus to induce snoRNA-jou degradation using shRNA interference. We used different amounts of lentivirus to infect the cells, corresponding to the multiplicity of infection (MOI). We first chose to study the effects on adenocarcinoma cells (HCT116), as we did on our previous studies [15], in order to see if siRNA-like effects were observed with this new sh-lentivirus. Second, to study the effects on primary cells, we chose glioblastoma primary cells GBM14. Cell proliferation of transduced cells was compared to non-transduced cells, to cells after application of polybren, and to a sh-scramble-lentivirus. Cell proliferation (viability) measurements were performed by two independent methods. First with the CellTiter-Glo assay, and second, directly counted by Malassez cells, at different times after transduction, as 72 hours, 8 and 10 days, to observe longer-term effects of shRNAs after the division of infected cells.

First, to determine the efficacy of the sh-lentivirus, we performed a dose-response effect of the cell proliferation (viability) on HCT116 cells measured at 72 hours post-treatment (Suppl. Fig. 1). The two controls, the polybren and the sh-control [mock (sh-scramble) lentivirus at a MOI20] did not generate any effect. However, we observe a nice decrease of cell proliferation by increasing the amount of sh-lentivirus anti-jouvence (MOI) used to infect cells. With a MOI-1, the cell proliferation is reduced by 10%, with MOI-5 by 20%, MOI-10 by 30%, MOI-20 by 50% and finally, MOI-50 by 60%. Then, we decide to perform the next experiments at a MOI-20, which will be compared to the sh-scramble control.

More specifically, at a MOI-20, at 72h post-transduction, HCT116 shows a decrease of 43% compared to sh-scramble, an effect confirmed by a Malassez counts (50% decrease) (Fig.1 A-B). To verify that the observed effects were not specific only to HCT116 cells, we transduced other immortalized cell types with sh-lentivirus. After 72 hours with MOI-20, we observed similar results on lung cancer cell line A549 with a reduced viability of 52% (Fig. 1C) and on glioblastoma cell line U87-MG with a reduced viability of 57% for MOI-20 (Fig. 1D). In parallel, to check molecularly the efficacy of the sh-lentivirus, RT-qPCR (TaqMan) were carried out in order to quantify the level of expression of snoRNA-jou in cells transduced by sh-lentivirus. With a MOI-20, at 72h, we found that transduced HCT116 cells underexpressed snoRNA-jou by 50% (fold change = 0,5) compared to untransduced cells (Fig. 1E).

**Figure 1.**
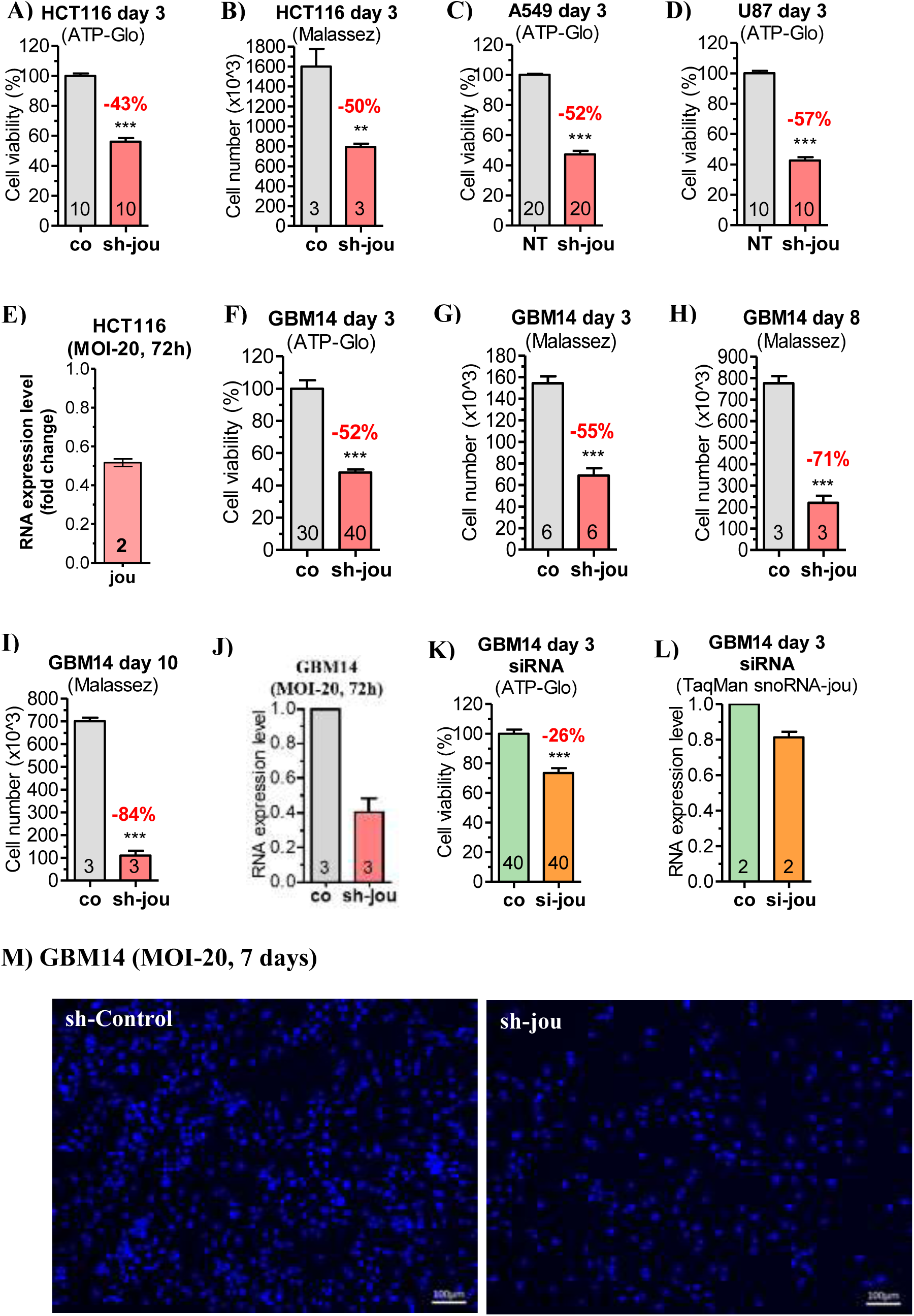
Knockdown of snoRNA-jou by sh-lentivirus decreases cancerous cell proliferation. **A-B)** HCT116 cells proliferation at 72 hours transduced with a sh-lentivirus control without target (sh-co), versus cells transduced with sh-lentivirus targeting snoRNA-jou at a MOI-20. **A)** Cell viability as determined by CellTiter-Glo, **B)** Number of cells as determined by Malassez cell counting. **C)** Lung cancer cell line A549 at 72 hours, MOI-20, as determined by CellTiter-Glo, **D)** Glioblastoma cell line U87-MG after 72 hours, MOI-20, as determined by CellTiter-Glo. **E)** In HCT116, the expression level of snoRNA-jou is decreased by about 50%, 72 hours after transduction (MOI-20). **F-G)** GBM14 cells proliferation at 72 hours transduced with a sh-lentivirus control without target (sh-co), versus cells transduced with sh-lentivirus targeting snoRNA-jou at a MOI-20. **F)** Cell viability as determined by CellTiter-Glo, **G)** Number of cells as determined by Malassez cell counting. **H)** as in G, but after 8 days post-transduction, **I)** after 10 days post-transduction. **J)** In GBM14, the expression level of snoRNA-jou is decreased by about 60%, 72 hours after transduction (MOI-20). **K)** GBM14 transduced with the siRNA, at 3 days post-transduction, as determined by the Cell-Titer-Glo. **L)** In GBM14, the expression level of snoRNA-jou is decreased by about 20%, 72 hours after transduction with siRNA. **M)** Microphotography of the GBM14 cells stained with Hoechst, 7 days post-transduction with sh-lentivirus compared to sh-scramble control. Statistics: (p-values : *** p < 0,0005). Errors bars represent the mean +/− SEM (p-value were calculated using the one-tail unpaired t-test using GraphPad Prism). Cell proliferation of each condition was compared to the sh-scramble (control) condition. For the CellTiter-Glo, numbers in each histogram represent the number of wells (e.g.: 40 = 4 time 10 wells, etc.), while for the Malassez counts, or the RNA expression level, it represents the number of replicates.

In a second step, as mentioned above, we investigate the sh-lentivirus on glioblastoma primary cells GBM14. Since primary GBM14 cells have twice the doubling time (about 60 hours) than the cells of immortalized line HCT116 (about 30 hours), the proliferation of the cells was measured at three different times post-transduction (72h, 8 and 10 days). After 72 hours, with the MOI-20, we observed a proliferation decrease of 52%, as measured by CellTiter-Glo (Fig. 1F), with a similar result (55%) when measured by direct Malassez cells count (Fig. 1G). After 8 days, we observe a decrease of 71% (Fig. 1H), and up to 84% at 10 days (Fig. 1I). As with HCT116, we checked at the molecular level, by RT-qPCR (TaqMan), the efficacy of the sh-lentivirus. With a MOI-20, we found that transduced GBM14 cells underexpressed snoRNA-jou by 60% (Fig. 1J). Obviously, in both cases, we observe a partial (not total) knockdown of the amount of snoRNA-jou. However, this partial inhibition is sufficient to lead to a striking effect on cell proliferation. We also check on the primary cancerous cells GBM14, as we did for the laboratory cell lines [15], if our previous siRNA (LNA type) were also efficient to decrease the cell proliferation. Indeed, Fig. 1K shows that the proliferation is reduced by 26% after 72h, which also roughly correspond to a reduction of about 20% of the amont of the snoRNA-jouvence (Fig. 1L). The difference in the % of reduction of cell proliferation observed between the siRNA and the sh-lentivirus is likely due the amount of siRNA entering into the cells, whose are also not integrated in the genome, and therefore lead to a more transient effect, compared to sh-lentivirus. Finally, the visual microscopic observations agree with cell proliferation analysis. Indeed, after 7 days post-transduction, the sh-control GBM14 shows much more cells than the sh-lentivirus treated cells (Fig. 1M). In addition, since the lentivirus used contains the eGFP, we take advantage of this property to evaluate the transduction efficiency. We could observe a transduction efficacy of almost 100% (see Suppl. Fig. 2). Altogether, these results confirm the efficacy of sh-lentivirus for the knockdown of the snoRNA-jou. In overall, these results therefore indicate that the knockdown of snoRNA-jou by sh-lentivirus induces a significant decrease in the cell proliferation of transduced cancer cells, with a dose-response effect, and with greater effects over time.

### EdU confirms a decrease of cell proliferation

To corroborate the decrease of cell proliferation revealed by the CellTiter-Glo and the direct determination of the number of cells by the Malassez count, we perform an EDU staining, which labels the cell in proliferation. As expected, for a MOI-20 at 72h, the number of labelled cells (in proliferation) treated with the sh-jou-lentivirus is reduced by about 30% compared to the cells treated with a sh-scramble lentivirus (Fig. 2A & B).

**Figure 2.**
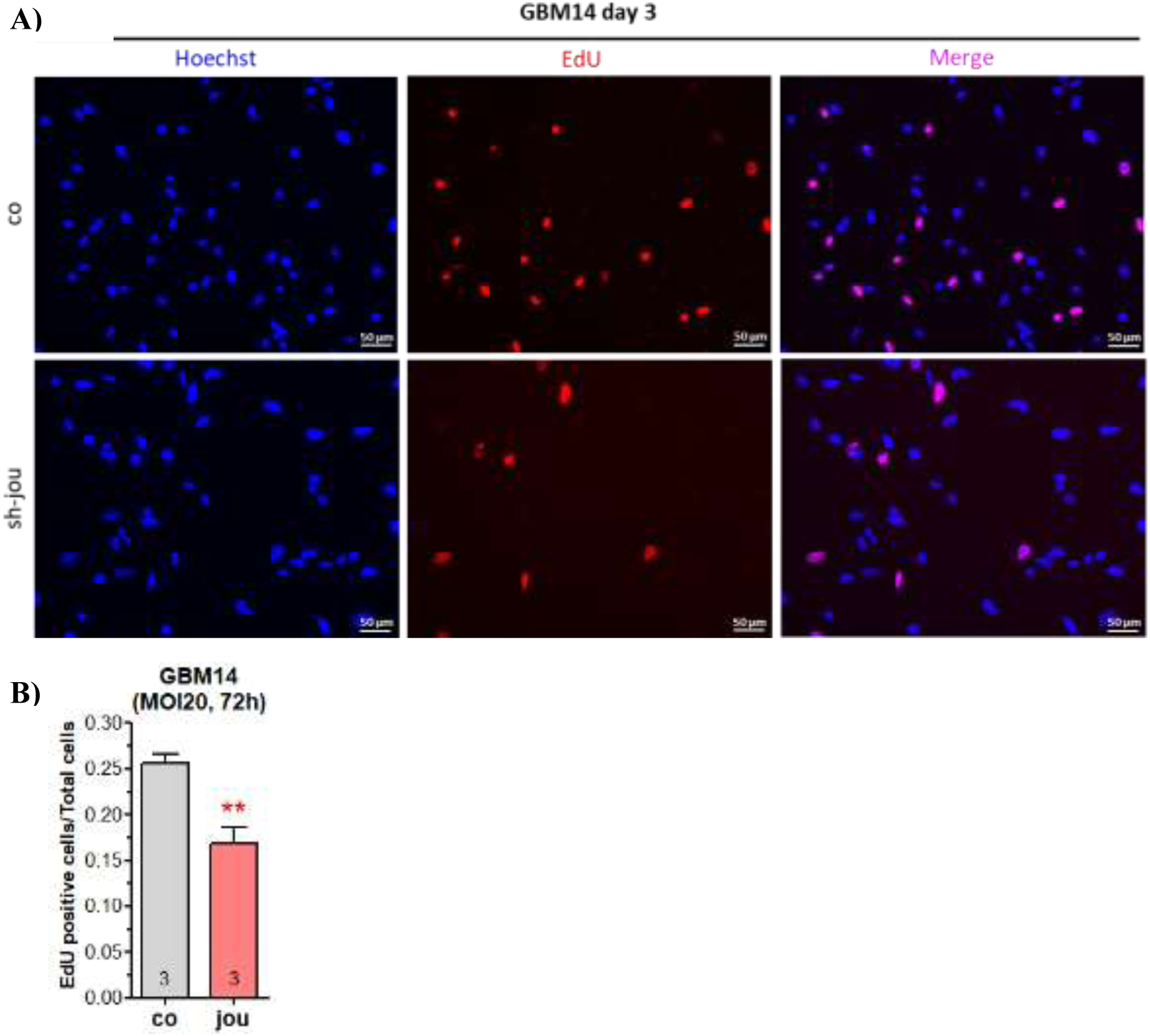
Knockdown of snoRNA-jou by sh-lentivirus reduces cell proliferation of GBM14 as revealed by EdU. **A)** Microphotography of the GBM14 cells treated wih sh-lentivirus compared to sh-scramble lentivirus (co), 3 days post-transdution (MOI-20). First column: Hoechst staining (in blue), Middle column: EdU staining (in red), Third column: Merge. **B)** Histogram showing the number of labelled EdU positive cells compared to total number of cells. (p-values : ** p < 0,001).

### TUNEL reveals the presence of apoptotic cells

In a step further, to asses if the sh-jou-lentivirus also leads to the death of the cells, we perform a TUNEL assay. For a MOI-20 at 72h, we remark that several cells are already in apoptosis (Fig. 3A & B), indicating that in addition to block the cell proliferation, several cells are entering in apoptosis and consequently dies.

**Figure 3.**
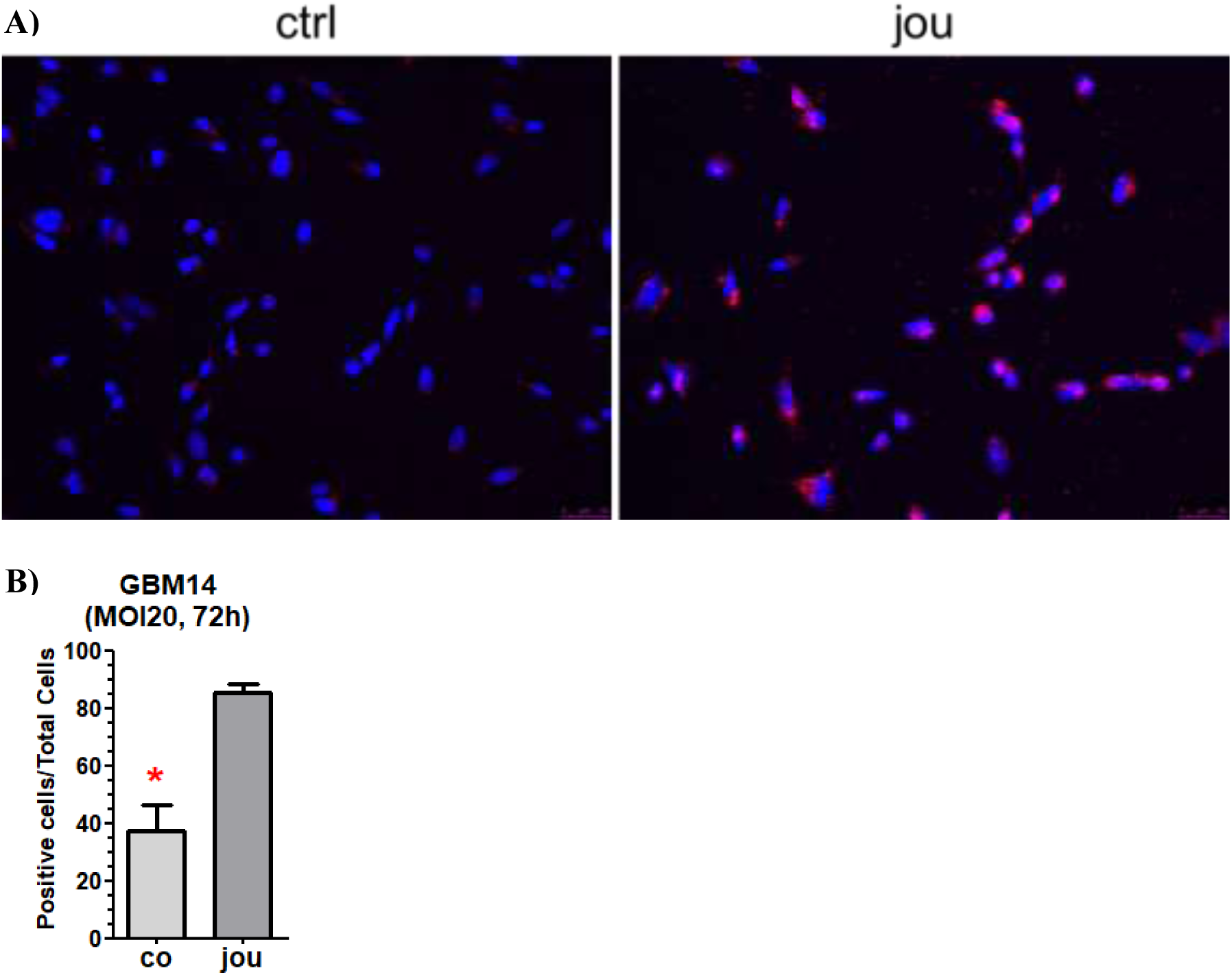
Knockdown of snoRNA-jou by sh-lentivirus induces cells death of GBM14 as revealed by TUNEL assay. **A)** Microphotography of the GBM14 cells treated wih sh-lentivirus compared to sh-scramble lentivirus (control-ctrl), 3 days post-transduction (MOI-20). In red: TUNEL staining, in blue: Hoechst staining. **B)** Histogram showing the number of labeled TUNEL positive cells compared to total number of cells. (p-values : * p < 0,05).

### Transcriptomic analysis (RNA-Seq) reveals cell cycle arrest

In order to elucide the molecular mechanisms responsible of the decrease of cell proliferation, we perform a transcriptomic analysis (RNA-Seq), by comparing the GBM14 treated with sh-jou-lentivirus (MOI-20, 72h) versus the GBM14 without treatment, as control. First, a Venn diagram reveals that 1254 expressed genes are specific of sh-lentivirus-treated cells, while 1033 are specific of control (Fig. 4). According to the standard criteria of a log2-Fold Change, and a p-value < 0,05, the differentially expressed genes (DEGs) analysis shows that 2953 genes are down-regulated, while 2743 genes are up-regulated (Fig. 5A) (see also the Suppl. Table 1 and 2 for the complete list of DEGs). However, more interestingly, the Gene Ontology (GO) analysis reveals that the cell cycle arrest is one of the most deregulated gene pathways (Fig. 5B) (see also the Suppl. Table 3 for the complete GO-enrichement). To point-out more precisely the deregulated genes involved, directly or indirectly in the cell cycle, the most important genes have been selected and gathered, based on the GO-enrichment (Table 1A & B). Notably, among the 56 down-regulated genes related to cell cycle (Table 1A) several cyclins and their related partners, as the cyclin dependent kinase inhibitor 2C (CDKN2C), cyclin dependent kinase inhibitor 3 (CDKN3C), cyclin dependent kinase inhibitor 2D (CDKN2D), cyclin dependent kinase inhibitor 1C (CDKN1C), cyclin dependent kinase 9 (CDK9), cyclin dependent kinase 5 (CDK5), cyclin B1 (CCNB1), and cell division cycle 25C (CDC25) (all highlighted in blue in Table 1A) are deregulated. Moreover, we also included in this Table some other deregulated cell cycle flagship genes, as growth arrest specific 1 (GAS1), GAS7, and GS2L1, as well as some more classical cancer flagship genes as tumor protein p53 (TP53), or p53-induced death domain protein 1 (PIDD). Similarly, among the 47 up-regulated genes (Table 1B), we also remark several cyclins, as cyclin D1 (CCND1), cyclin dependent kinase inhibitor 2A (CDKN2A), cyclin dependent kinase 7 (CDK7), or the cell division cycle 14A (CDC14A). Moreover, several other cyclins are also up-deregulated (highlighted in green in Table 1B), as cyclin L1, cyclin J, cyclin Y, cyclin B1-interacting protein 1 (CCNB1IP1), cyclin H, cyclin Y like 1, cyclin T2, and cyclin E2, although these last have not been indexed directly in the cell cycle in the GO analysis. Finally, the KEGG analysis did not reveal striking difference, only few pathways are statistically downregulated, while none pathway are up-regulated (see Suppl. Table 4 for the complete KEGG analysis pathways).

**Figure 4.**
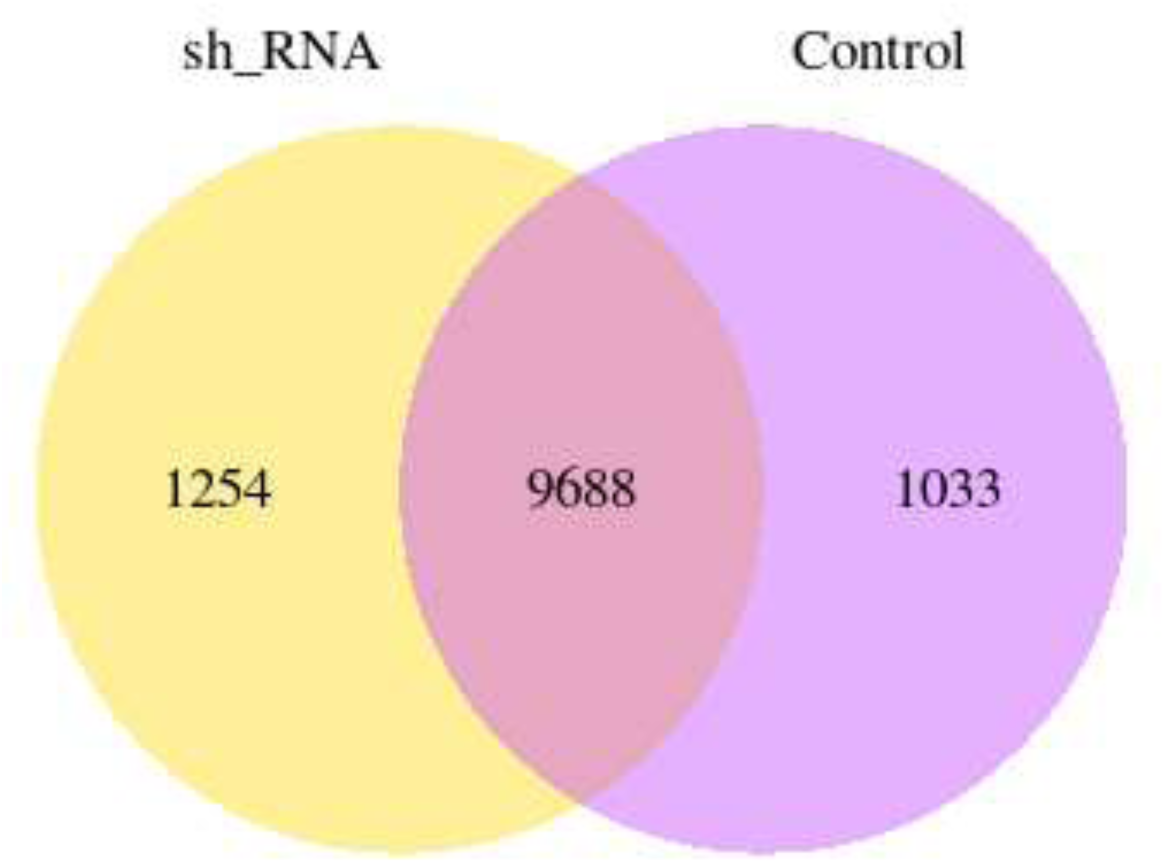
Venn diagram of the transcriptomic analysis comparing GBM14 sh-lentivirus treated cells with control untreated GBM14. 9688 genes are expressed in common, while 1033 are expressed only in control cells, and 1254 only in sh-lentivirus treated cells.

**Figure 5.**
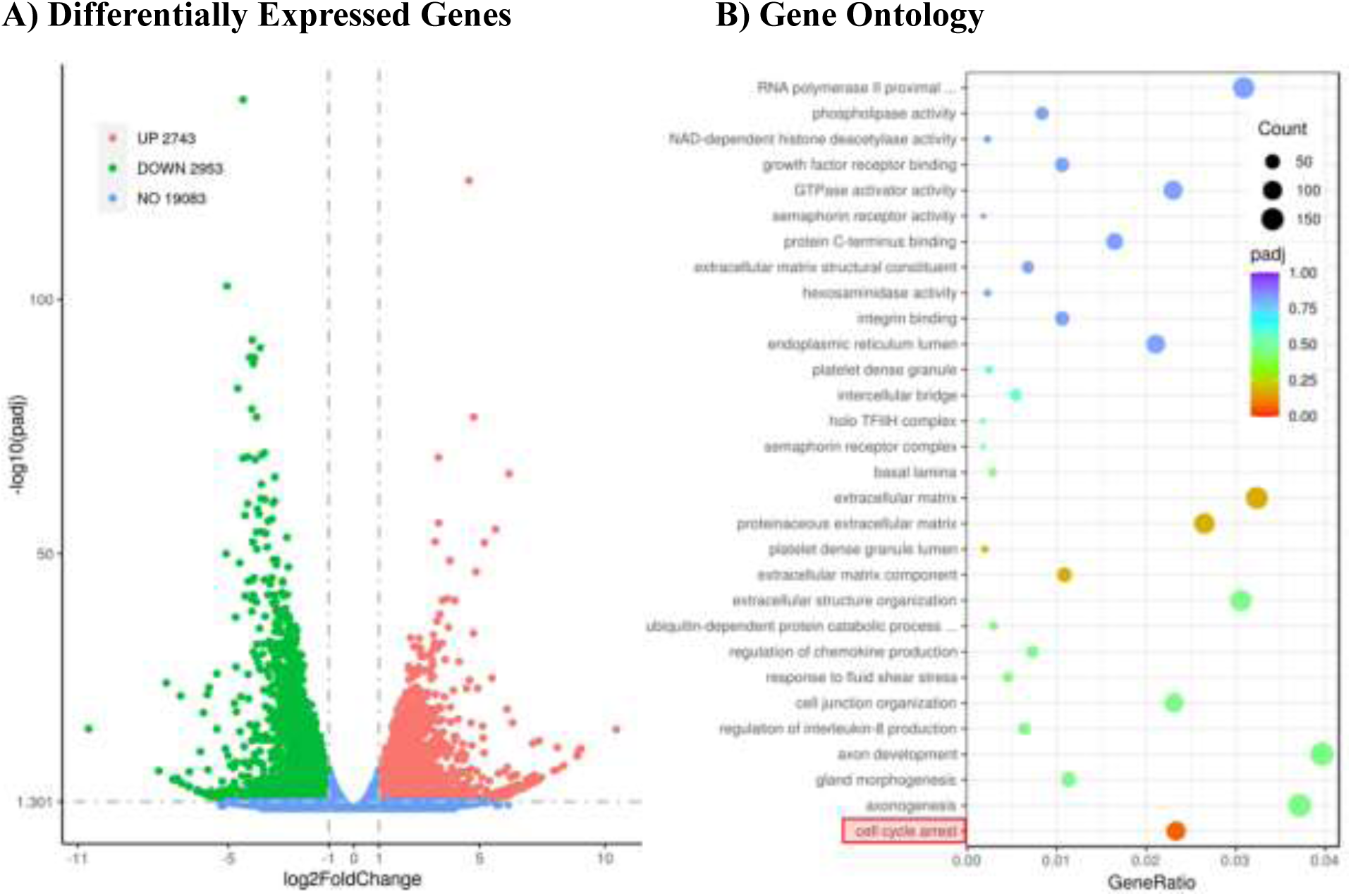
Transcriptomic analysis performed on total-RNA from the knockdown of snoRNA-jou by sh-lentivirus on GBM14 cells compared to the control non-transduced cells. **A)** Differentially Expressed Genes (DEGs) reveals that 2743 genes are upregulated, while 2953 are downregulated (see Suppl. Table-S1 and S2 for the complete list of genes). **B)** Gene Ontology analysis (GO) revealing several deregulated pathways, and notably the cell cycle arrest (see Suppl. Table-S3 for the complete list of GO-enrichment).

**Table 1:**
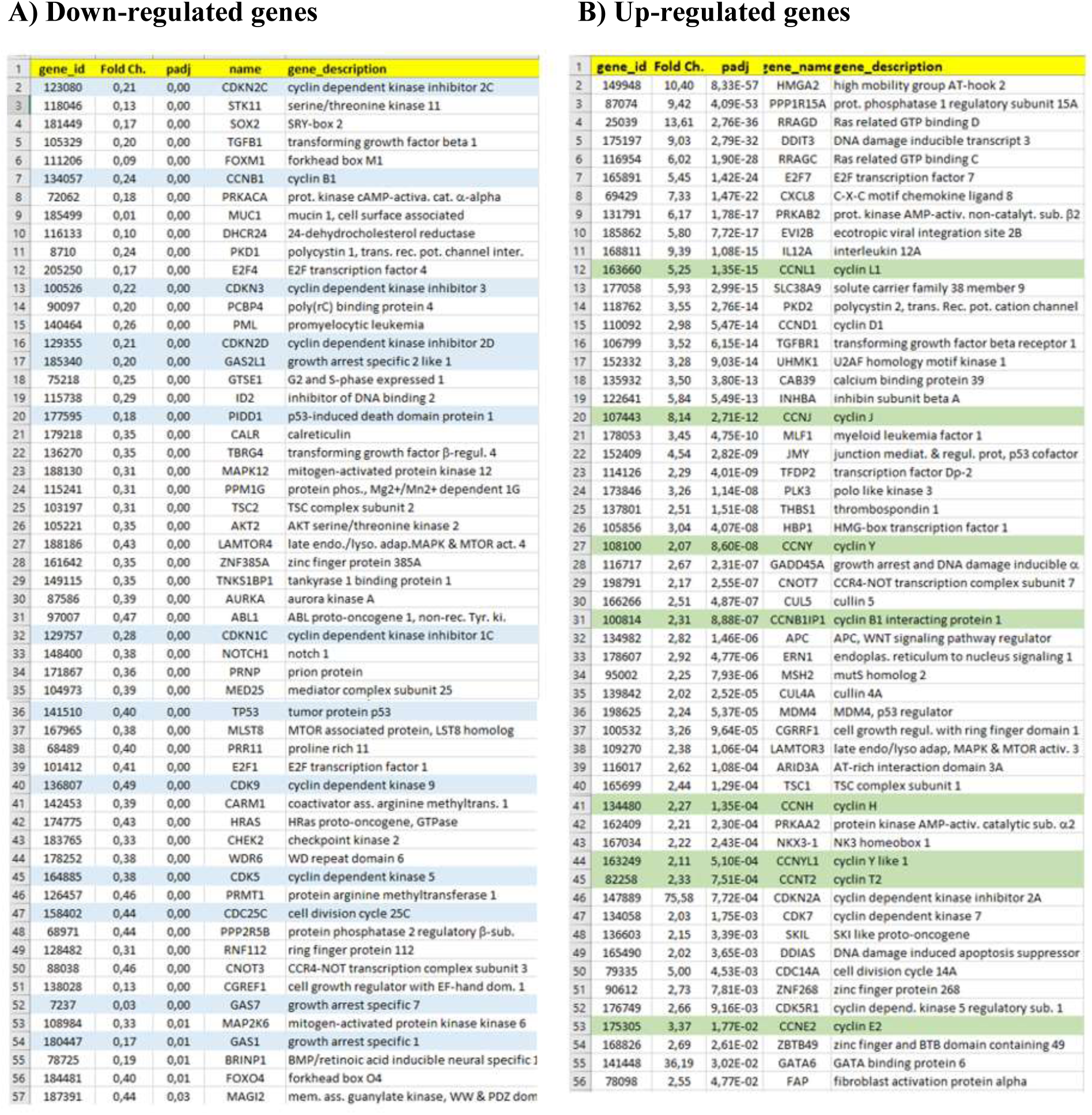
List of Deregulated Genes (up and down) involved in the cell cycle revealed by the GO-analysis. Some additional genes, not repertoried in the GO analysis, but also known to be related to the cell cycle have been included (highlighted in blue in A, and green in B).

### RT-qPCR validation of the dysregulation of several genes involved in cell cycle

In order to characterize more precisely cellular and molecular mechanisms affected by the knockdown of snoRNA-jou, and in the aim to validate them by a second independent approach, analysis of the expression of several interesting deregulated genes was performed by RT-qPCR. We selected genes that were the most deregulated in the RNA-Seq analysis performed both on HCT116 cells treated with siRNAs directed against snoRNA-jou [15], and on the GBM14 cells (cited above). In the colon adenocarcinoma cells HCT116, knockdown of snoRNA-jou by sh-lentivirus induces a down-regulation of the MKI67 (proliferation marker), CCNB1 (encoding the cyclin B1 protein), CCNA2 (encoding the cyclin A2 protein), CCND1 (encoding the cyclin D1 protein) MYC, RPLPO, TP53, HDAC5, PRPF3 and SNRNP70 genes (Fig. 6A). Knockdown of snoRNA-jou also induces significant overexpression of CDKN1A (encoding the p21 protein), SLC7A11 and HMGCR genes (Fig. 6A). On primary glioblastoma cells GBM14, knockdown of snoRNA-jou induces a down-regulation of MKI67, TP53, CCNB1 and CCNA2 genes (Fig. 6B). Overexpressions of the CDKN1A, CCND1 (encoding the cyclin D1 protein), MYC, HMGCR and SLC7A11 genes are also observed (Fig. 6B). To summarize, in these two independent cell types, the commun underexpressed genes are the MKI67, cyclin B1 (CCNB1), cyclin A2 (CCNA2) and TP53 genes, while the overexpressed genes are the p21 (CDKN1A), SLC7A11 and HMGCR genes. Interestingly, the cyclin B1, A2, MKI67 and p21 genes encode for proteins directly involved in cell cycle regulation. Indeed, the p21 protein (overexpressed here) is an inhibitor of cyclin-cdk complexes (cyclins B1 and A2 are underexpressed here). These results therefore suggest an effect of snoRNA-jou knockdown on the cell cycle of sh-lentivirus anti-jou transduced cells.

**Figure 6.**
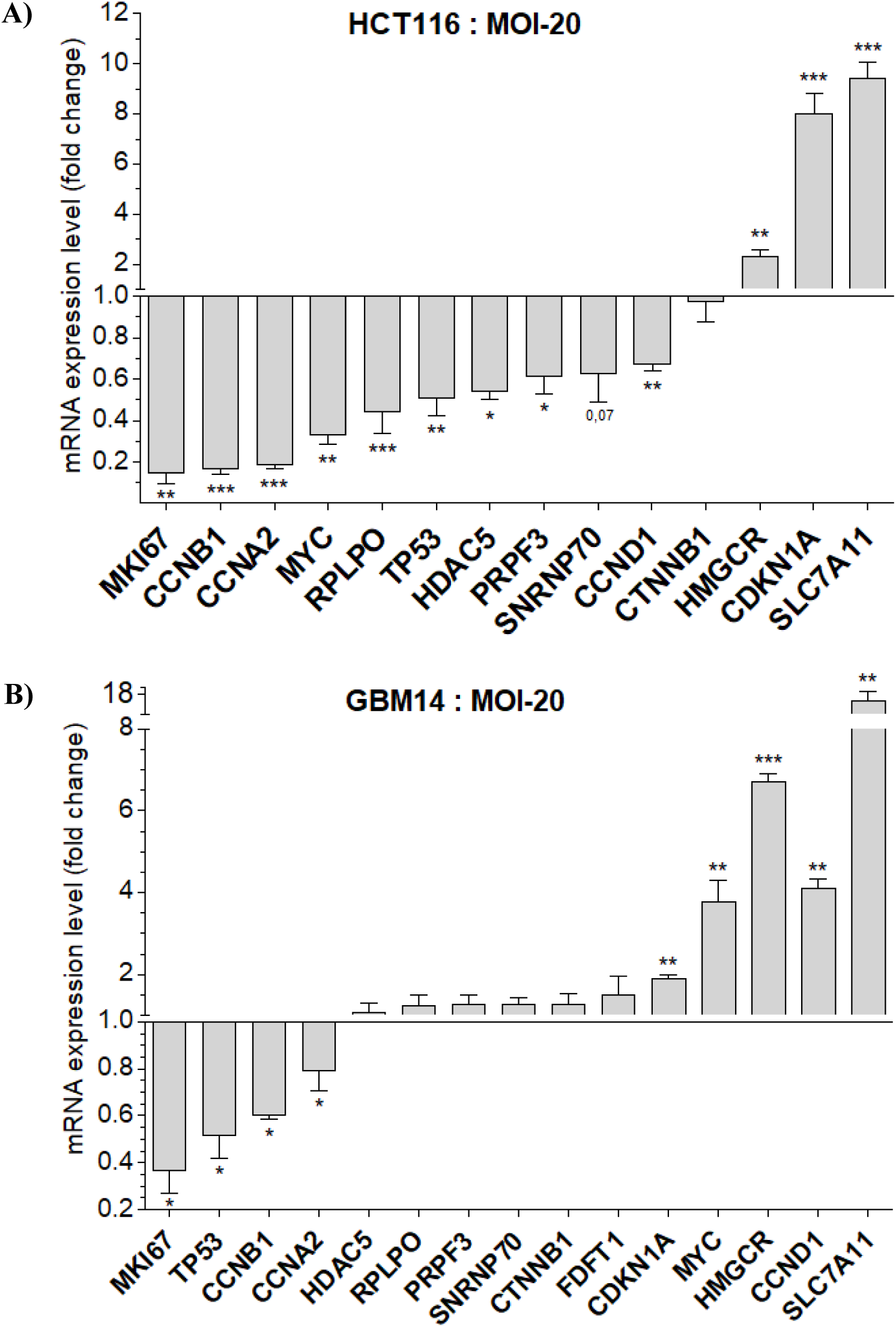
RT-qPCR validation of the knockdown of snoRNA-jou by sh-lentivirus inducing dysregulation of several genes. Dysregulated genes in **A)** HCT116 and **B)** GBM14 cells after transduction by sh-lentivirus compared to non-transduced cells. The genes are classified according to their level of deregulation. RNAs were extracted from cells harvested 72 hours after MOI-20 sh-lentivirus transduction. Gene expression is normalized by the housekeeping gene GAPDH expression. Statistics: Two independent experiments have been performed for each figures (n=2). Errors bars represent the mean +/− S.E.M. p-value were calculated using the one-tail paired t-test on the delta-CT values using GraphPad Prism.

To confirm that the mRNA deregulation of the genes leads also to the deregulation of the protein, we also perform Western Blots on some proteins. Then, Western blots were performed to quantify the expression of the p21 protein in HCT116 cells at 72 hours after transduction by sh-lentivirus at MOI-20. We observed that the p21 protein was 4 times more expressed in transduced cells compared to non-transduced cells (Fig. 7A & B). Moreover, we also demonstrated the underexpression of the cyclin B1 protein in transduced cells compared to non-transduced cells (Fig. 7C & D). These results therefore confirm at the protein level the dysregulation of the p21 (up-regulated) and cyclin B1 (down-regulated) genes observed by RT-qPCR after the knockdown of the snoRNA-jou.

**Figure 7.**
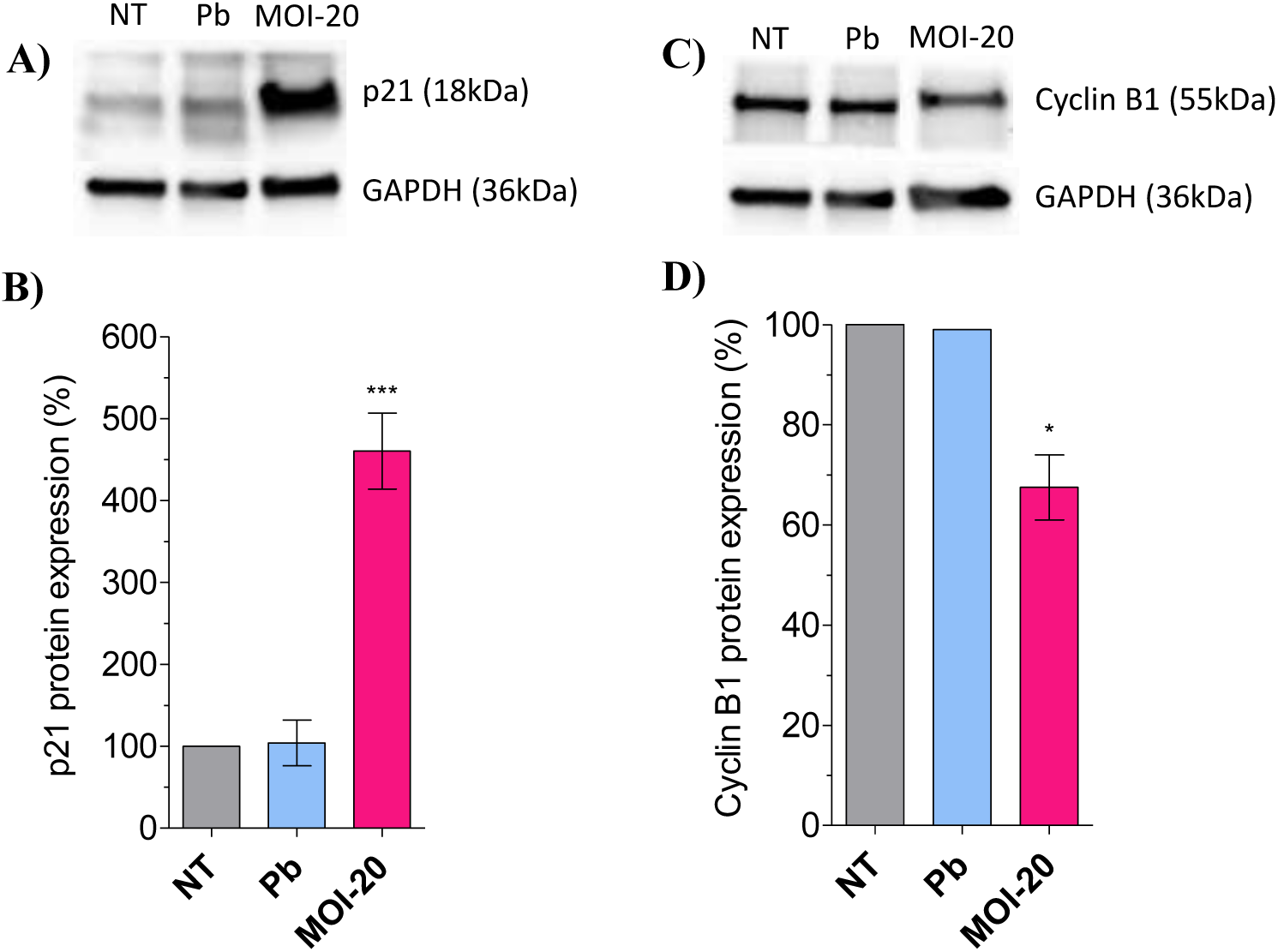
Western Blots of the p21 and cyclin B1 protein expression levels following the knockdown of snoRNA-jou by sh-lentivirus. **A)** p21 and GAPDH proteins gel electrophoresis. **B)** Relative quantification of p21 protein expression. **C)** Cyclin B1 and GAPDH proteins gel electrophoresis. **D)** Relative quantification of cyclin B1 protein expression. Results obtained on HCT116 cells with polybrene application (Pb) and in cells transduced with sh-lentivirus (MOI-20) compared to non-transduced cells (NT). Statistics : p-values : * p < 0,05 ; *** p < 0,0005. Errors bars represent the mean +/− S.E.M. (p-value were calculated using the one-tail unpaired t-test using GraphPad Prism).

### Lipidomic Analysis reveals a deregulation of Cholesterol

Cholesterol homeostasis has been implicated in several cellular and molecular mechanisms, and notably in cancer [17,18], while some snoRNAs have already been involved in cholesterol metabolism [19]. As revealed by the transcriptomic analysis, several key genes involved in cholesterol pathways are deregulated, among them the 3-Hydroxy-3Methyl-Glutaryl-Coenzyme A Reductase (HMGCR) (Fig. 6), the limiting enzyme of the cholesterol synthesis. Thus, to investigate if the cholesterol is indeed deregulated, as well as other lipids, we performed a lipidomic analysis on GBM14 cells treated with the sh-lentivirus anti-jouvence. At a MOI-20 and 72h post-transduction, we observe a strong increase in various form of cholesterol, as cholesterol itself, the dihydrocholesterol and the cholesterol esters (Fig. 8). We also observe an increase in the total lipids, the triglycerides and the total phospholipids (Suppl. Fig. 3). However, the precise determination of the composition of the fatty-acid (FA) of these various classes of lipids did not reveal striking difference between the two conditions (sh-scramble-control versus sh-lentivirus anti-jou) (Suppl. Fig. 4). Indeed, in overall, in all lipid classes, each FA is increased by about a factor 3 in the sh-lentivirus anti-jou treated cells compared to their controls, roughly similar to the increase observed for the corresponding lipid class itself. Altogether, these results indicate that the knockdown of jouvence perturbs quantitatively the overall and general lipid and cholesterol homeostasis, without affecting specifically a precise FA pathway.

**Figure 8.**
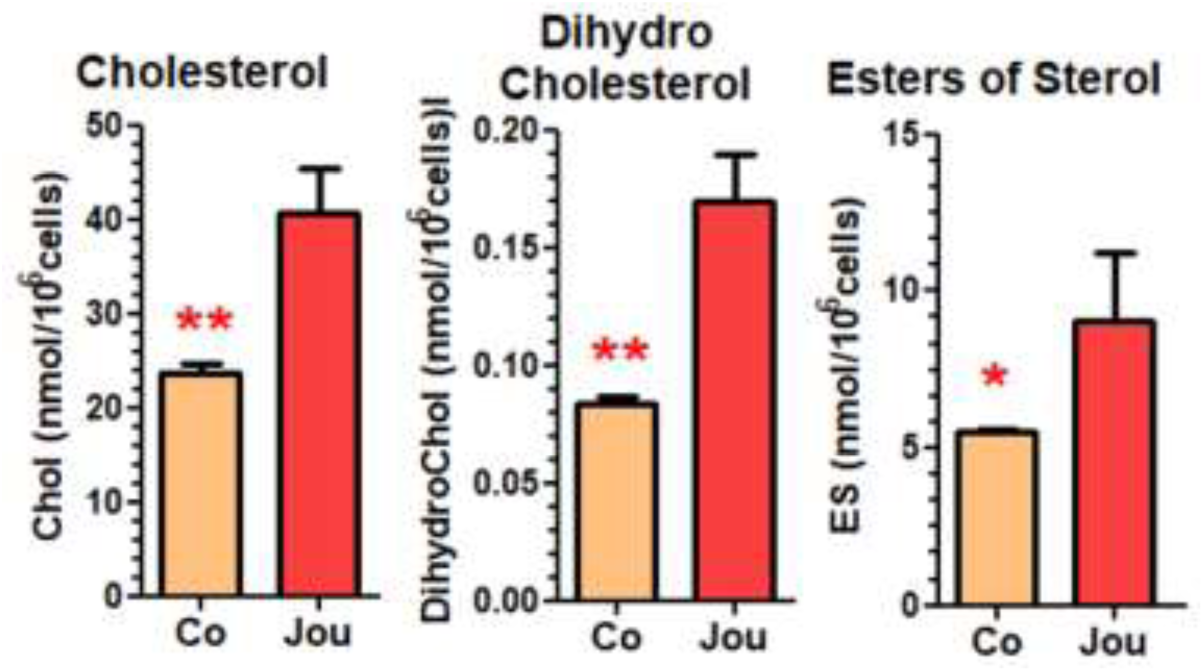
Lipidomic analysis reveals an important increase in different forms of Cholesterol. Knockdown of snoRNA-jou by sh-lentivirus compared to sh-scramble lentivirus (co), 3 days post-transduction (MOI-20), induces an important increase in Cholesterol, DihydroCholesterol and Esters of Sterol. Statistics : p-values : * p < 0,05 ; ** p < 0,005. Errors bars represent the mean +/− S.E.M. (p-value were calculated using the one-tail unpaired t-test using GraphPad Prism) (n=3 replicates).

## DISCUSSION

This study validates the use of lentiviruses containing shRNAs as a tool for inhibition (knockdown) of snoRNA-jou, both on three immortalized cell lines, and more importantly, on patient-derived primary (glioblastoma) cells (xenograft), a terrifying cancer known to be refractory to almost all types of treatment. Indeed, on the colon adenocarcinoma cell line (HCT116), the lung cancer cell line A549, and on glioblastoma cell line U87-MG, as well as on primary glioblastoma cells (GBM14), transduction by sh-lentiviruses induces a significant decrease in cell proliferation, and as demonstrated in HCT116 cells, proportional to the amount of lentivirus used (from MOI-1 up to MOI-50) (Suppl. Figure 1). With a MOI of 20, a decrease of more than 50% in cell proliferation is observed after only 72 hours in colon adenocarcinoma cells (HCT116), as well as in primary glioblastoma cells. The integration of shRNA-jou into the cell genome seems to be functionally validated by the analysis done over longer periods. Indeed, cell proliferation decreases more significantly over time, demonstrating long-term effects and therefore a probable presence of shRNA-jou in the daughter cells after the division of the infected cells. This result is also supported by si-RNA transfection, although the observed effect is weaker in this last. In addition, the downregulation of snoRNA-jou was demonstrated, by RT-qPCR, in treated cells, thus validating the use of sh-lentivirus as a knockdown tool for snoRNA-jou. Finally, as revealed by the RNA-Seq analysis, the knockdown of snoRNA-jou induces significant deregulation of several genes, and particularly including several key genes involved in cell cycle regulation. Interestingly, in GBM14 cells, the Venn Diagram (Fig. 4) has revealed that 1033 genes are expressed only in the control line, meaning that their expression is totally extinct in the treated cells. In contrary, 1254 genes are newly expressed in treated cells, while they were not expressed in non-treated controls cells. These results suggest, likely indirectly, a role of the snoRNA-jou in the regulation (positive or negative) of transcription.

As we have confirmed and displayed in the Fig. 6, several genes are communly deregulated in both studied cell types (HCT116 and GBM14). More particularly, the genes encoding cyclin B1 (CCNB1) and cyclin A2 (CCNA2) exhibited a strong decreased expression in cells treated with shRNA-jou. The cell cycle is a fundamental process allowing the production, from a mother cell, of two identical daughter cells. Cyclin/cdk complexes allow the transition between the different phases of the cell cycle of eukaryotic cells. These complexes are composed of a regulatory subunit, cyclin, and a catalytic subunit, cyclin-dependent kinase (cdk) (Fig. 9). Cyclins have a cyclic expression, thus conditioning and regulating the activation of cdk at certain times of the cell cycle. The cyclin B1/cdk1 complex regulates the G2/M transition of the cell cycle to allow entry into mitosis; while cyclin A2 associates with cdk2 or cdk1 to regulate the S and G2 phases [20,21]. In this study, the underexpression of cyclins B1 and A2 in cells treated with shRNA-jou strongly suggests a decrease in the activation of cyclin/cdk complexes and therefore a blockage of the cell cycle.

**Figure 9.**
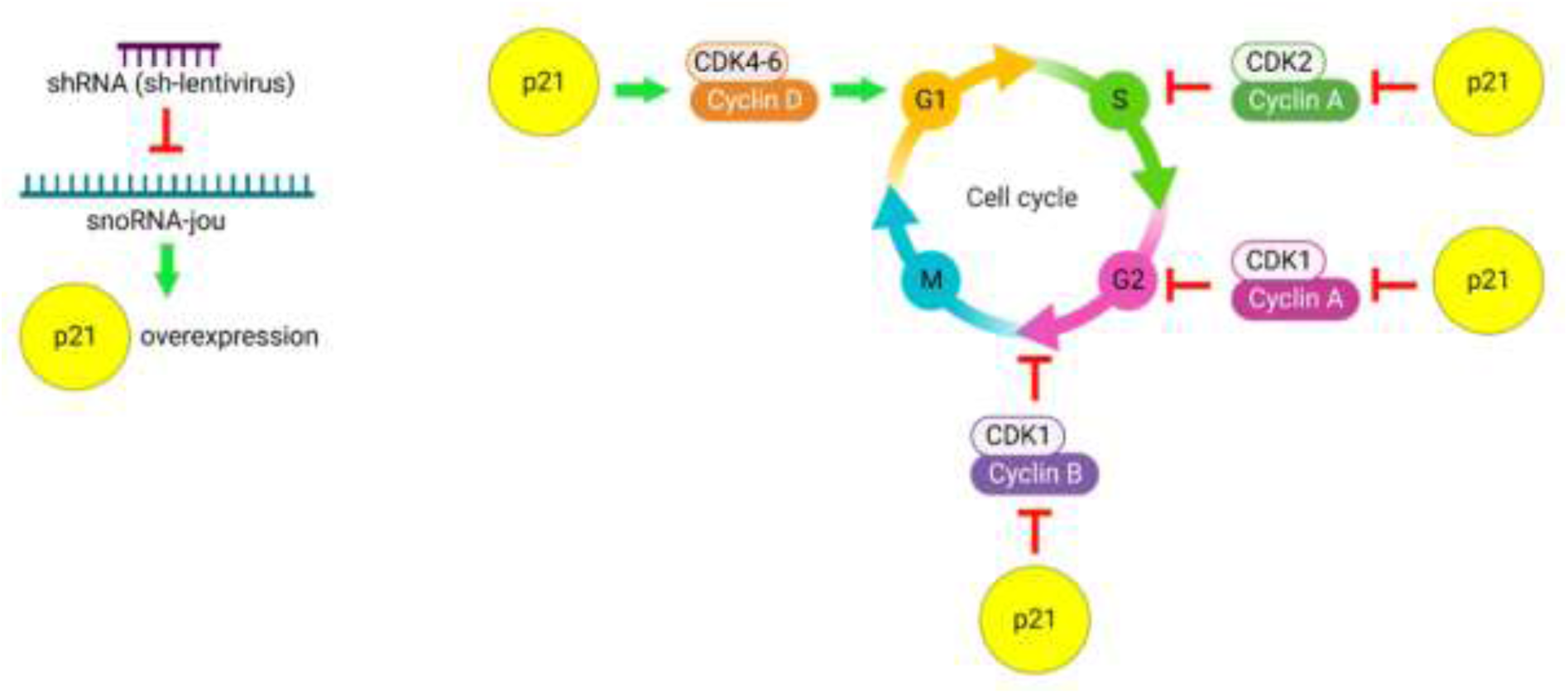
Working model depicting how the snoRNA-jou knockdown might affect the cell cycle. Knockdown of snoRNA-jou by sh-lentivirus induces overexpression of p21 protein. p21 protein inhibits the cyclin A-CDK2 complex thereby inhibiting the S phase. p21 protein inhibits also the cyclin A-CDK1 complex thereby inhibiting the G2 phase. p21 protein also inhibits the cyclin B-CDK1 complex thereby inhibiting G2/M transition.

The p21 protein is an inhibitor of the cyclin E/cdk2 complex, thus disrupting the checkpoint of the G1/S transition (Fig. 9) and inducing a blockage of the cell cycle in the G1 phase [21]. Knockdown of snoRNA-jou induces overexpression of the p21 protein, supporting also here, the hypothesis of a cell cycle blockage, probably in G1 phase. The Ki-67 marker (MKI67) is a proliferation marker whose expression varies according to the phases of the cell cycle. Indeed, the Ki-67 marker is weakly expressed during the G0 and G1 phases, and strongly expressed during the S, G2, and M phases [22]. In the treated cells, the expression of the Ki-67 marker is drastically reduced, supporting the hypothesis of a cell cycle blockage in the G1 phase. The MYC oncogene, encoding the transcription factor c-Myc, which positively regulates cell growth and proliferation, contributes to the genesis of many human cancers [10,11]. Indeed, studies have shown that an aberrant expression of c-Myc promotes aerobic glycolysis, resulting in providing a significant amount of energy for the initiation, maintenance and growth of certain tumors [23]. In vivo studies in a mouse model of lung adenocarcinoma have also demonstrated that systemic inhibition of c-Myc allows rapid regression of incipient and established tumors [24]. Here, in accordance with our finding, in HCT116 cells, snoRNA-jou knockdown induces significant underexpression of the MYC oncogene, suggesting a positive effect in inhibiting cancer cell growth.

The p53, a transcription factor, also well known as a tumor suppressor gene, which has been involved in many cancers [25] is down-regulated both in HCT116 and in GBM14 sh-lentivirus treated cells, compared to their non-treated cells respectively. Alhough this decrease does not seem, at first glance, compatible with a decrease of the proliferation of cancerous cells, we have to keep in mind that here, whatever the case may be, several other genes are deregulated, and that the outcome of this complex deregulation, which remain to be thoroughly elucidated, leads to a decrease of cell proliferation.

It is also interesting to observe that several other genes, as for instance, involved in pre-mRNA splicing are strongly deregulated after snoRNA-jou knockdown. Indeed, the PRPF3 (Pre-mRNA Processing Factor 3), known to associate to the snRNP U4 and U6 for the elimination of introns from the pre-mRNA, and SNRNP70 gene (Small Nuclear Ribonucleoprotein U1 Subunit 70), which enables U1 snRNA binding activity, and which is involved in mRNA splicing, via spliceosome and regulation of RNA splicing, are under-expressed in HCT116 cells. A disruption of splicing mechanisms could have consequences on the proliferation of cancer cells by altering the structure of mRNAs and translated proteins [26].

As mentioned above, cholesterol homeostasis has been implicated in several cellular and molecular mechanisms, as well as in pathophysiology of certain cancers [17,18]. Here, in sh-lentivirus anti-jou treated GBM14 cells, the HMGCR gene, the rate-limiting step in cholesterol biosynthesis, is up-regulated, which correlates to an elevated level of different forms of cholesterol (cholesterol, dihydro-cholesterol and cholesterol esters), as determined by a lipidomics analysis. Cholesterol and its metabolites have already been involved in different cancers [17,18,27], and/or associated to apoptosis [28–30]. Interestingly, it has been reported that up-regulation of cholesterol synthesis results from p53 functional loss [31], a correlation also observed here between the p53 and HMGCR. However, if the increase of HMGCR level is a direct effect of the p53 decrease remains to be determined. Obviously, as already underlined by other studies, a relationship between the cholesterol level, the apoptopsis and the cell cycle is also observed here.

To conclude, we have demonstrated that the knockdown of a small non-coding RNA, the snoRNA-jou, has a significant anti-proliferative effect on cancer cells, including patient-derived primary cells, probably associated with a deregulation of the cell cycle. Chemotherapy drugs are designed to inhibit different mechanisms of cancer cells, such as DNA replication, protein synthesis or tumor angiogenesis. Recent studies suggest that chemotherapy combined with cell cycle regulation strategies have positive effects for cancer treatment [21]. Although snoRNAs do not code for proteins, they have demonstrated an important involvement in many cellular processes such as cell cycle regulation. Increasing studies are currently pointing small nucleolar RNAs as promising potential biomarkers of cancer, either for detection or prognostic signature [2,32,33]. Therefore, targeting non-coding RNAs, such as the snoRNA-jou, represents an original and promising new strategy for cancer therapy.

## METHODS

### Cell lines and culture conditions

#### Immortalized cell lines

HCT116 (human colon adecarcinoma), A549 (human lung carcinoma) and U87-MG (human glioblastoma) cell lines comes from the American cell bank ATCC (American Type Culture Collection (USA).

#### GBM14-CHA primary cells

The patient-derived primary GBM14-CHA cells (abbreviated GBM14) come from the company XenTech (Evry-Courcouronnes, France), which specializes in the production of PDX (Patient-Derived Xenograft) cells. PDX cells are human tumor cells grafted into immunodeficient mice in order to be amplified, then dissociated and cultured. These are human glioblastoma cells taken from a 70-year-old woman.

#### Culture conditions

Cells are cultured according to standard protocols. HCT116 cells are cultured in McCoy’s 5A medium (Gibco™) supplemented with 10% fetal calf serum (Gibco™). A549 cells are cultured in RPM1 medium (Gibco™) supplemented with 10% fetal calf serum (Gibco™). U87-MG cells are cultured in DMEM+GlutaMAX medium (Gibco™) supplemented with 10% fetal calf serum (Gibco™). GBM14 cells are cultured in DMEM/F-12+GlutaMAX medium (Gibco™) supplemented with 8% fetal calf serum (Gibco™) and 1% penicillin-streptomycin (Gibco™). Cells were incubated at 37°C and 5% CO2.

### Lentivirus transduction

The ultra-purified lentivirus were designed by ourself, and were produced by VectorBuilder™. They allow the expression of a short hairpin RNA (shRNA) targeting the sequence of snoRNA-jou (GGCTATTGTGGACAGAGGA) or a negative (CCTAAGGTTAAGTCGCCCTCG) control sequence (sh-scrambled). The viral particle titer of the lentivirus tubes is 3.10^9^ TU/ml. On the first day, the cells are counted, seeded and incubated for 24 hours. The next day, the lentivirus are applied to the cells in different quantities. Mixes containing medium, polybrene (increases transduction efficiency) and lentivirus are made according to the multiplicity of infection (MOI) determined. The MOI corresponds to the ratio between the number of viral particles used to infect the cells and the number of cells. The culture medium in which the cells were seeded is removed, then the mixes containing the lentivirus are added to each well. The following day, the medium containing the lentivirus is removed and replaced with fresh medium. The control condition is cells treated with control “scramble” lentivirus (shRNA without target in the cell) abbreviated “co”.

### CellTiter-Glo cell viability assay

The CellTiter-Glo test (Promega™) measures cell viability, based on the quantification of ATP, an indicator of metabolically active cells. The results obtained correspond to a quantity of luminescence, considered to be proportional to the number of living cells. The measurements are carried out on 96-well plates with a white background. CellTiter-Glo reagent is added directly to the wells containing the cells. The plate is shaken away from light (15 min), then the luminescence is measured with the GloMax Navigator plate reader (Promega™). For a one-plate read, each condition is represented by 10 wells. “Blank” wells with medium without cells are also measured. The analysis of the results consists of subtracting the average of the blanks from each raw data, then dividing these values by the average of the control condition (untreated cells) and multiplying them by 100. The average of each condition is thus calculated to obtain a percentage of relative luminescence, compared to the control condition. HCT116, A549 and U87-MG cells are seeded at 3000 cells and GBM14 cells are seeded at 5000 cells per well.

### Cell Counting

Cells are counted using standard Malassez counting chamber.

### Transcriptomic analysis (RNA-seq)

The transcriptomic analysis (RNA-seq) was performed by Novogene (China, the Cambridge U.K. Branch). Briefly, total RNA was extracted and quality control performed. Libraries enriched for polyA RNAs, were generated and checked for quality. The libraries were sequenced on an Illumina Hiseq platform, and 125 bp/150 bp paired-end reads were generated. For all the computational tools analysis, common defaults parameter values were used. The reference genome and gene model annotation files were downloaded from genome website directly. Index of the reference genome was built using Bowtie v2.2.3 and paired-end clean reads were aligned to the reference genome using TopHat v2.0.12. We selected TopHat as the mapping tool because TopHat can generate a database of splice junctions based on the gene model annotation file and thus a better mapping result than other non-splice mapping tools. For the quantification of gene expression level, HTSeq v0.6.1 was used to count the read numbers mapped to each gene. Then FPKM of each gene was calculated based on the length of the gene and read counts mapped to this gene. FPKM, expected number of Fragments Per Kilobase of transcript sequence per Millions base pairs sequenced, considers the effect of sequencing depth and gene length for the read counts at the same time, and is currently the most commonly used method for estimating gene expression levels.

Differential expression analysis of two conditions (three biological replicates per condition) was performed using the DESeq R package (1.18.0). DESeq provide statistical routines for determining differential expression in digital gene expression data using a model based on the negative binomial distribution. The P values were adjusted using the Benjamini & Hochberg method. Corrected P-value of 0,05 and log2 (Fold-change) of 1 were set as the threshold for significantly differential expression. The biological variation was eliminated (case with biological replicates), and the threshold was therefore normally set as p-adjusted (padj) <0,05. Gene Ontology (GO) enrichment analysis of differentially expressed genes was implemented by the GOseq R package, in which gene length bias was corrected. GO terms with corrected p-value less than 0,05 were considered significantly enriched by differential expressed genes. For the KEGG database [34,35], KOBAS software was used to test the statistical enrichment of differential expression genes in KEGG pathways.

### RT-qPCR

RNAs are extracted from the cell pellets using the NucleoSpin RNA Plus kit (Macherey-Nagel™). DNAse treatment is performed with RQ1 RNAse-free DNAse (Promega™). RNAs are then precipitated with ethanol and 0.3M sodium acetate (−20°C, over-night), centrifuged (4°C, 30 min) and resuspended in RNAse-free water. Reverse transcription reactions are performed with M-MLV Reverse Transcriptase (Invitrogen™) with random primers (qPCR TaqMan) or oligo dT (qPCR SYBR Green). The template RNA strands are then removed by treatment with RNAse H (Thermo Scientific™). qPCR reactions are performed using Master Mix PowerUp SYBR Green (Applied Biosystems™) and TaqMan Universal Master Mix II (Applied Biosystems™). The Taqman probe is the same than the one used in El-Khoury et al., (2020)[15]. For the Primers sequences, see Suppl. Table 5.

### Western blot

Extraction of total proteins, dosage of protein lysates, migration, transfer and revelation are carried out according to standard protocols. The membrane is incubated with the primary antibody solution at 4°C overnight with shaking. Primary antibodies directed against the p21 protein (SC6246, Santa Cruz Biotechnology) are diluted 1:200. Primary antibodies directed against the cyclin B1 protein (SC53236, Santa Cruz Biotechnology) are diluted 1:2000. Primary antibodies directed against the GAPDH protein (2118S, Cell Signaling Technology) are diluted 1:1000. The membrane is then incubated with the secondary antibody solution for two hours at room temperature. For cyclin B1, anti-mouse HRP secondary antibodies (P0447, Agilent) are diluted 1:5000. For p21 and GAPDH proteins, anti-rabbit HRP secondary antibodies (SC7074, Santa Cruz Biotechnology) are diluted 1:2000.

### Cell proliferation Assay (EdU)

Proliferation tests were carried out using the Click-iT EdU assay with Alexa Fluor 594 (Invitrogen), according to the supplier’s instructions. Cells were seeded in 8-wells Lab-tek (Thermo Scientific). GBM14 cells are seeded at 10000 cells per well.

### Apoptotic cells detection (TUNEL)

Apoptotic cells detection was carried out using the Elabscience® One-step TUNEL In Situ Apoptosis Kit according to the supplier’s instructions. Cells were seeded in 8-wells Lab-tek (Thermo Scientific). GBM14 cells are seeded at 10000 cells per well.

### Lipidomic Analysis

Lipid analysis was performed on the Functional Lipidomics Platform acknowledged by IBiSA (Infrastructure in Biology, Health and Agronomy). After addition of appropriate internal standards (1,2-diheptadecanoyl-*sn*-glycero-3-phosphocholine, cholesteryl ester 17:0, tri-17:0 triglyceride and ^13^C cholesterol,) to the samples, total lipids were extracted twice with chloroform/ethanol (2:1, v/v). The organic phases were dried under nitrogen and the different lipid classes were then separated by Thin Layer Chromatography using the solvent mixture hexane-diethylether-acetic acid (80:20:1, v/v/v) as eluent. Lipids were revealed by UV light after spraying the plate with 0.02% dichlorofluorescein in ethanol and identified by comparison with standards. Silica gel was scraped off. Total lipids, total phospholipids, triacylglycerols and cholesteryl esters were transmethylated and the fatty acid methylesters were analyzed by gas chromatography. Briefly, samples were treated with toluene-methanol (2:3, v/v) and 14% boron trifluoride in methanol. Transmethylation was carried out at 100°C for 90 min. The reaction was stopped by cooling the samples to 0°C and by the addition of 1.5 ml K_2_CO_3_ in 10% water. The resulting fatty acid methyl esters were extracted with 2 ml of isooctane and analyzed by gas chromatography using an HP6890 instrument equipped with a fused silica capillary BPX70 SGE column (60 × 0.25 mm), with hydrogen as a vector gas. Temperatures of the flame ionization detector and the split/splitless injector were set at 250°C and 230°C, respectively. Peak detection and integration were performed using ChemStation software.

Cholesterol was extracted twice with chloroform/ethanol (2:1, v/v). The dry residue was derivatized with 100 μl N,O-bis-trimethylsilyl-trifluoroacetamide (BSTFA) for 20 min at 60°C. Derivatized cholesterol was then analyzed by gas chromatography (HP 7890B, Agilent) coupled with triple quad mass spectrometry (GC-MS/MS) using the electron impact ionization (EI) mode (7000C, Agilent). GC-MS/MS was equipped with a SolGel-1ms fused silica capillary column, 60 m length, 0.25 mm internal diameter (Trajan, SGE). The carrier gas was helium with a constant flow rate of 1.2 ml/min. The split/splitless injector was heated to 280°C. The oven temperature program started at 55°C for 4 min and then ramped up to 250°C at 40°C/min, and finally up to 310°C at 20°C/min for 25 min. The samples were injected in a splitless mode. The temperature of the mass spectrometer transfer line was set at 250°C, while the source temperature was set at 200°C. Nitrogen was used as collision gas (1.5 ml/min). The electron ionization energy was 70 eV. Cholesterol was detected using the multiple-reaction monitoring (MRM) mode.

### Statistical analysis

Data were analysed statistically using one-tailed unpaired t-test with GraphPad Prism software.

## Supporting information

Supplementary Table-5 (Primers)

## Acknowledgements

We thank Juliette Bitard (NeuroPSI) for her excellent advices and assistance in culture cells. This work was supported by the Fondation ARC pour la Recherche sur le Cancer (ARCPJA32020060002114), and the ANR (Aging-jou) to JRM, as well as by the CNRS to J. Bignon (JBi) and JR Martin. The authors declare that they have no competing interests.

## Author contributions

JRM conceived and designed the experiments. LJC, JBu, and NBH performed experiments. LJC, JBu, JBi, NBH, and JRM analysed the data. LJC wrote the first draft of the manuscript, and JRM wrote the final version of the manuscript with input from all authors.

## Conflict of interest

The authors declare no conflict of interests.

**Supplementary Information:** Four Supplementary Figures, and five Supplementary Tables (S1 to S4) containing the RNA-Seq Data-analysis (Excel files), and Table (S5) containing the sequence of Primers used for the RT-qPCR, are available.

## Data Availability

The authors declare that all the data and the methods used in this study are available within this article, its Supplementary Information files, or are available from the corresponding authors upon reasonable request.

