## Supplementary Figures 1-4 for "The knockdown of the snoRNA-jouvence blocks the cell proliferation and leads to cell death of human primary cancerous glioblastoma cells"

**Suppl. Fig. 1) Dose response effect of different MOI of the sh-lentivirus transduction.**

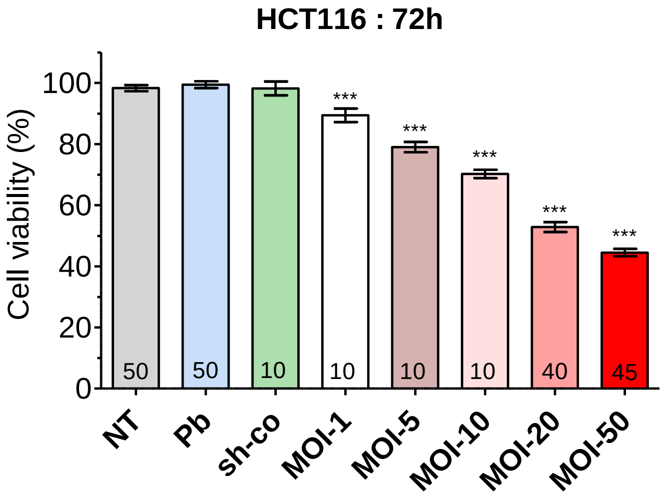


**Suppl. Fig. 2) Efficiency of the sh-lentivirus transduction (MOI-20, 72h post-transduction)**

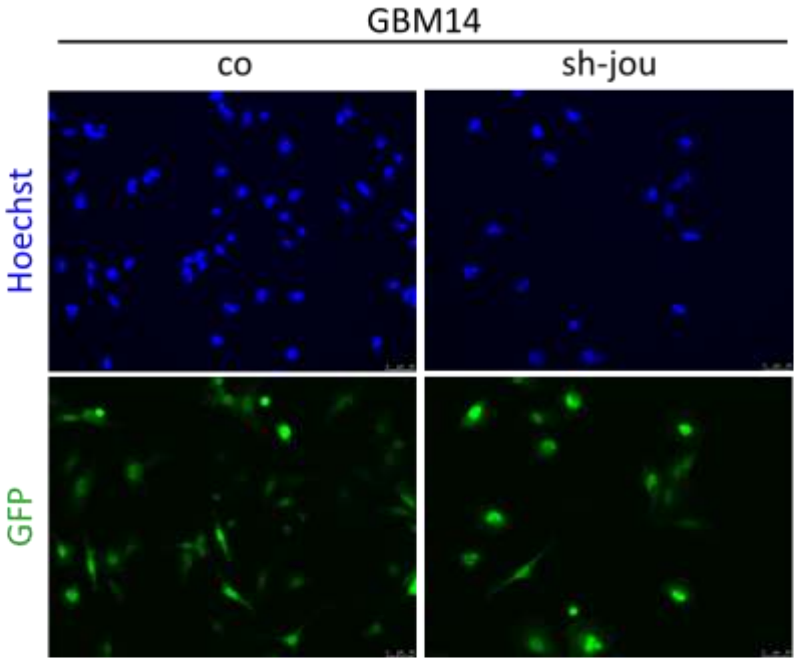

**Suppl. Fig. 3) Total amount of different classes of Lipids.**

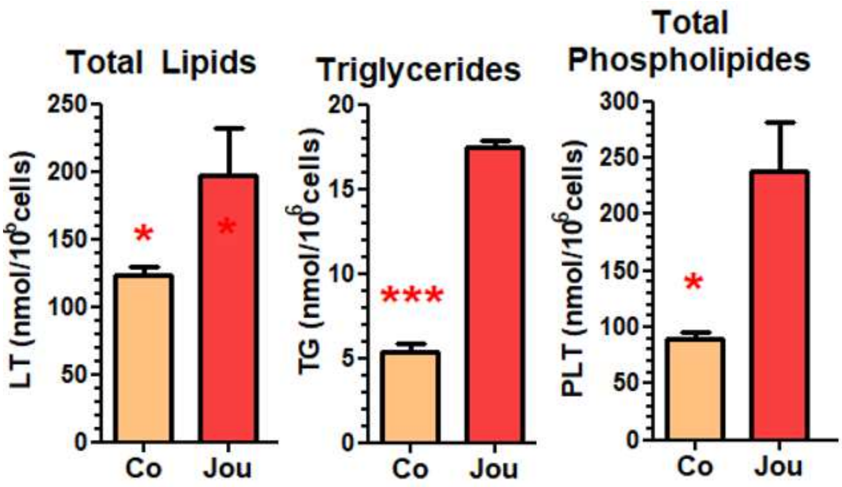

**Suppl. Fig. 4) Fatty-acid composition of the different classes of Lipids and Cholesterol** (from data of Fig. 8 and Suppl. Fig. 3). FA= Fatty-Acid, FA-S = FA-Saturated, FA-MU = FA-Mono-Unsaturated, FA-PU = FA-Poly-Unsaturated, FA-PU-w3 = FA-Poly-Unsaturated omega-3, FA-PU-w6 = FA-Poly-Unsaturated omega-6, w3/w6 = ratio omega-3/omega-6.

### A) Fatty Acids in Total Lipids (LT)

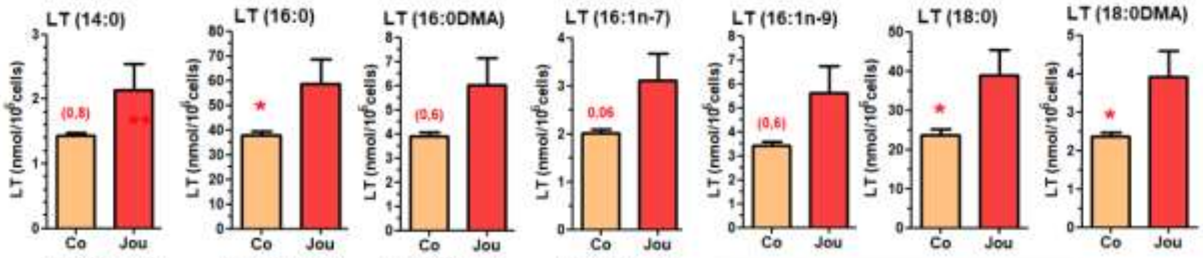


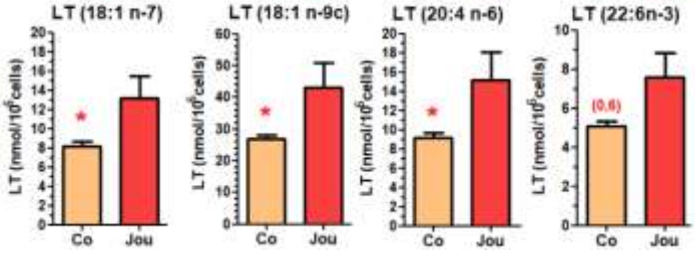


| FA | Co | Jou |
|---|---|---|
| FA-S | 69,27 | 109,60 |
| FA-MU | 40,36 | 64,97 |
| FA-PU | 9,14 | 15,12 |
| FA-PU-w3 | 5,09 | 7,57 |
| FA-PUw6 | 9,14 | 15,12 |
| w3/w6 | 0,56 | 0,50 |

### B) Fatty Acids in Triglycerides (TG)

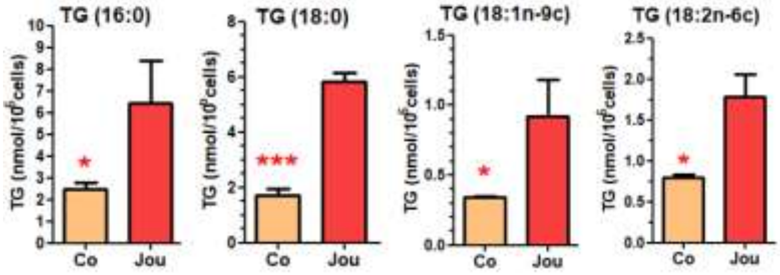


| FA | Con | Jou |
|---|---|---|
| FA-S | 4,19 | 10,91 |
| FA-MU | 0,34 | 0,92 |
| FA-PU | 0,79 | 1,79 |
| FA-PUw3 | 0,00 | 0,00 |
| FA-PUw6 | 0,00 | 0,00 |
| w3/w6 | 0,00 | 0,00 |

### C) Fatty Acids in Esters of Sterols (ES)

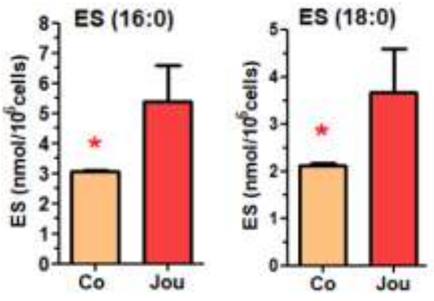


| FA | Con | Jou |
|---|---|---|
| FA-S | 5,18 | 6,77 |
| FA-MU | 0,35 | 0,00 |
| FA-PU | 0,00 | 0,00 |
| FA-PUw3 | 0,00 | 0,00 |
| FA-PUw6 | 0,00 | 0,00 |
| w3/w6 | 0,00 | 0,00 |

**D) Fatty Acids in Total Phospholipids (PLT)**

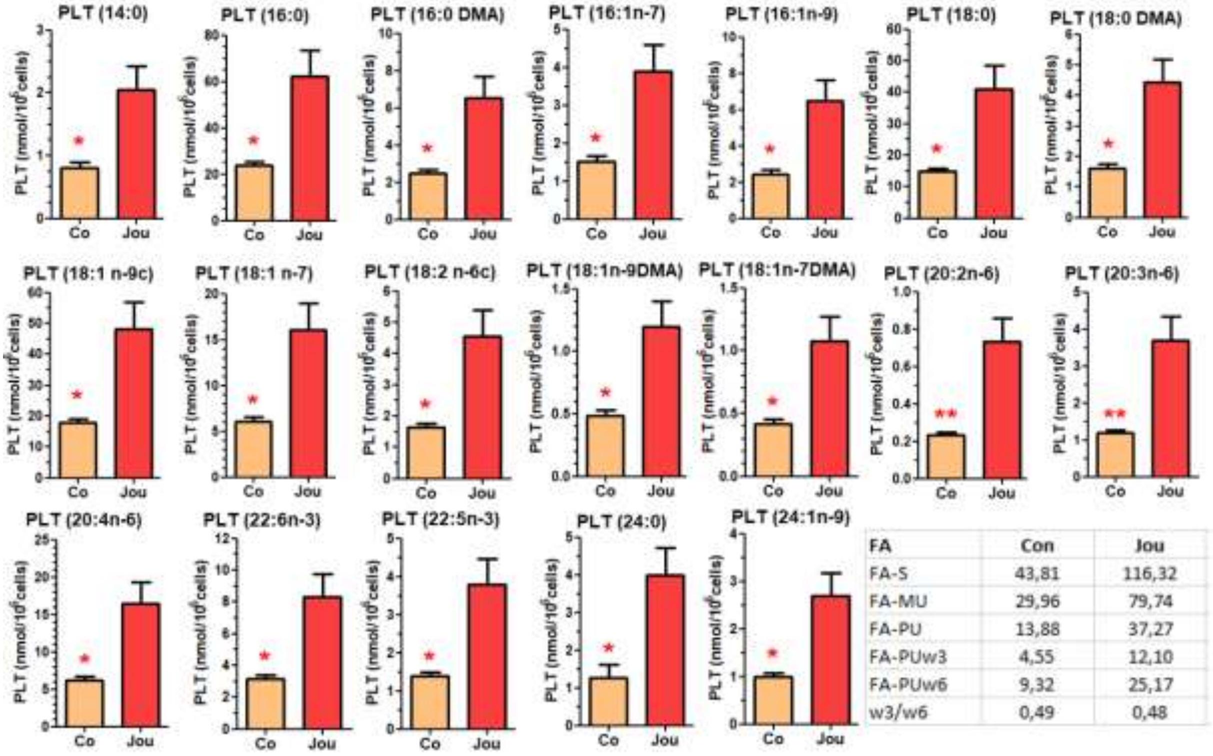


| FA | Con | Jou |
|---|---|---|
| FA-S | 43,81 | 116,32 |
| FA-MU | 29,96 | 79,74 |
| FA-PU | 13,88 | 37,27 |
| FA-PUw3 | 4,55 | 12,10 |
| FA-PUw6 | 9,32 | 25,17 |
| w3/w6 | 0,49 | 0,48 |
