## Supplementary Table-5 (Primers) for "The knockdown of the snoRNA-jouvence blocks the cell proliferation and leads to cell death of human primary cancerous glioblastoma cells"

| Gene | Forward primer | Reverse primer |
|---|---|---|
| CCNA2 | TTATTGCTGGAGCTGCCTTT | CTCTGGTGGGTTGAGGAGAG |
| CCNB1 | CCTGCTGCAACCTCCAAG | TCAGGTTCTGGCTCAGGTTC |
| CCND1 | AGAAGCGAGAGCCGAGC | TTGAAGTAGGACACCGAGGG |
| CDKN1A | GTGGACCTGTCACTGTCTTGTAC | CTTCCTCTTGGAGAAGATCAGC |
| CTNNB1 | GAAACGGCTTTCAGTTGAGC | CTGGCCATATCCACCAGAGT |
| FDFT1 | GGTCCCGCTGTTACACAACT | AAAACTCTGCCATCCCAATG |
| GAPDH | GAGTCAACGGATTTGGTCGT | TTGATTTTGGAGGGATCTCG |
| HDAC5 | GTGACACCGTGTGGAATGAG | AGTCCACGATGAGGACCTTG |
| HMGCR | GTCATTCCAGCCAAGGTTGT | TCCTGTCCACAGGCAATGTA |
| MKI67 | AGCCCCAACCAAAAGAAAGT | GACCTACGGCGTTGATCACT |
| MYC | GGGGACACTTCCCCGCCGCTG | CAGTAGAAATACGGCTGCACCGAG |
| PRPF3 | CAAGGAGGGAAGCACAGAAG | GCTCTGACGTGGGCTTCTAC |
| RPLP0 | GCGACCTGGAAGTCCAACTA | TCTCCAGAGCTGGGTTGTTT |
| SLC7A11 | GGCAGTGACCTTTTCTGAGC | TCATTGTCAAAGGGTGCAAA |
| SNRNP70 | CAGTAAGCGGTCAGGAAAGC | ACATCAGCCCCTCCTCTTCT |
| TP53 | GCCCAACAACACCAGCTCCT | CCTGGGCATCCTTGAGTTCC |

Legend: Suppl. Table-5 :  Sequences of the primers used in this study.
